# Murine limb ischaemia induces structural and functional remodelling of local and distant bone marrow

**DOI:** 10.1101/2025.09.16.676622

**Authors:** Lauren Eades, Michael Drozd, Nadira Y Yuldasheva, Anna Skromna, Natasha Makava, Nicole Powell, Jessica Smith, Oliver I Brown, Natalie J Haywood, Alexander F Bruns, Matthew C Gage, Marc A Bailey, Kathryn J Griffin, Stephen B Wheatcroft, Mark T Kearney, Richard M Cubbon

## Abstract

Peripheral arterial disease (PAD) can induce bone marrow (BM) ischaemia, although little is known about the implications of this for local and systemic haematopoiesis. We explored the impact of murine unilateral hind-limb ischaemia (HLI) surgery on the structure and function of the ischaemic and non-ischaemic BM, versus control mice without HLI surgery. Abnormal BM hypoxia was present in the ischaemic limb 7-days post-HLI, with normalisation by 28-days. Histological analysis revealed the ischaemic BM to undergo profound vascular remodelling at 7-days that normalised by 61-days, along with a progressive accumulation of BM lipid droplets between 7- and 61-days; no structural changes were observed in the non-ischaemic limb BM. Flow cytometry revealed increased abundance of monocytes and inflammatory monocytes in the non-ischaemic and ischaemic BM 7-days post-HLI, with this persisting at 61-days in the non-ischaemic limb. In both limbs, we observed a progressive decline in the abundance of lineage^−^Sca-1^+^c-Kit^+^ haematopoietic stem cells. RNA-sequencing of BM-derived macrophages (BMDMs) revealed substantial transcriptomic differences between unstimulated control, ischaemic limb and non-ischaemic limb BMDMs. Moreover, there were also substantial transcriptomic differences between lipopolysaccharide-stimulated control and ischaemic limb BMDMs. However, we observed minimal differences in chromatin accessibility of unstimulated BMDMs from the 3 groups. Collectively, our data show that BM ischaemia has transient and sustained structural implications, along with both local and systemic effects on haematopoiesis and myeloid cell function. These findings suggest that PAD may have important consequences for systemic immune responses.

## Introduction

Peripheral arterial disease (PAD) describes the presence of atherosclerosis in the arterial system outside the heart and brain and most commonly affecting the lower extremities. It affects over 200 million people across the world and is linked to reduced quality of life and life expectancy.(1) Whilst this is asymptomatic in a significant proportion, classically it causes exertional symptoms related to inadequate limb perfusion i.e. claudication. Around one in ten people with PAD experience chronic limb-threatening ischaemia (CLTI), characterised by ischaemic rest pain, tissue loss, or gangrene lasting at least 2 weeks. In keeping with its atherosclerotic basis, PAD is more prevalent with advancing age, and is closely associated with smoking, diabetes and dyslipidaemia. Indeed, it is also frequently associated with atherosclerosis in other vascular beds, especially the coronary artery and cerebrovascular disease.

Whilst the cutaneous and soft-tissue implications of CLTI are evident to patients and clinicians through symptoms and signs of tissue ischaemia, the impact of bone marrow (BM) ischaemia receives little attention in clinical practice. This is an important oversight, since limb BM is a source of haemopoiesis, with locally produced leukocytes, platelets and erythrocytes being released into the systemic circulation. Whilst only a proportion of haematopoiesis occurs in the lower limbs, the example of clonal haematopoiesis of indeterminate potential (CHIP) illustrates the capacity of even a small subset of altered leukocytes to promote a wide range of inflammatory processes.(2)

Only a few pre-clinical studies have explored the impact of BM ischaemia, all using the murine hind-limb ischaemia (HLI) model. The work of Falero-Diaz *et al* has shown that 24-hours of hind-limb ischaemia-reperfusion injury leads to a sustained change in BM-monocytes from the ischaemic limb as assessed at the transcriptional, lipidomic and functional level.(3) Notably, these cells were more effective in promoting neovascularisation of ischaemic muscle, with follow on studies showing priming of BM-macrophages to an anti-inflammatory M2-like phenotype.(4) However, Coppin *et al* demonstrated that sustained HLI in atherosclerosis prone ApoE^−/−^ mice promotes pro-inflammatory myeloid cell egress from the BM, which was associated with increased atherogenesis.(5) Using single cell RNA-sequencing (RNA-seq), they found perturbations in inflammatory gene expression in all myeloid cell populations, along with altered gene expression in their progenitors and haematopoietic stem cells (HSCs).

These data highlight the impact of HLI on BM function, but many important uncertainties remain. First, it is unknown how HLI impacts on BM structure and composition, both in the acute and chronic phases. Second, the impact of HLI on immune cell responses to inflammatory factors has not been described. Third, it is unclear whether only the ischaemic limb BM is altered, or whether there is a systemic impact on BM. To address these gaps in the literature, we characterised ischaemic and non-ischaemic limb BM at multiple timepoints after HLI, contrasting with control littermate mice not undergoing HLI. We hypothesised that HLI would alter BM structure and macrophage responses to lipopolysaccharide (LPS), both in the ischaemic and non-ischaemic BM.

## Methods

### Murine husbandry and ethics

C57BL/6J mice were purchased from Charles River Laboratories (UK) and used at 8-10 weeks of age. Mice were maintained under humidity- and temperature-controlled conditions with a 12-hour light/dark cycle and fed a standard chow diet *ad libitum*. Both male and female mice were used in all experiments. All experimental procedures were approved by the Animal Welfare and Ethical Review Body at the University of Leeds and conducted in accordance with The Animals (Scientific Procedures) Act 1986 Amendment Regulations 2012 under United Kingdom Home Office project licences P144DD0D6 and PP2103311.

### HLI surgery

HLI surgery was performed as previously outlined.(6) Mice were anaesthetised with isoflurane and the left femoral vessels dissected. The femoral artery was ligated proximal to the saphenous artery bifurcation and distal to the inguinal ligament and popliteal vessels, and the intervening arterial segment excised. Sham surgery was performed on the contralateral (right) hindlimb involving femoral vessel dissection but without ligation or excision. Buprenorphine (0.1 mg/kg in saline) was administered subcutaneously for analgesia.

### Laser Doppler imaging

Laser Doppler perfusion imaging was performed under isoflurane anaesthesia on hindlimbs using the Moor LDI2-HR system (Moor systems, UK) in a temperature-controlled environment. Perfusion was quantified using MoorLDI software (v6, Moor systems, UK) by calculating the relative perfusion in the ischaemic hindlimb compared to the contralateral hindlimb based on flux below the level of the knee joint.

### BM histology and confocal microscopy

#### Tissue preparation

BM histology was performed as previously described by Kusumbe *et al*.(7) Mice were euthanised, and femurs and tibiae harvested. Bones were fixed in 4% paraformaldehyde in PBS for 4 hours at 4°C and washed in PBS. Bones were decalcified in 0.5M EDTA/PBS (pH 7.4–7.6) for 24 hours at 4°C with agitation. After PBS washes, bones were cryoprotected in 20% sucrose/2% polyvinylpyrrolidone (PVP) in PBS for 24 hours at 4°C, then embedded in 8% gelatin/20% sucrose/2% PVP in PBS at 60°C, set for 1 hour at room temperature (RT) and stored at −80°C. Longitudinal bone sections (50μm) were cut using a Leica CM3050S cryostat.

#### Hypoxia assessment (*Hypoxyprobe*^*TM*^)

To evaluate BM oxygenation, mice were intraperitoneally injected with 120 mg/kg pimonidazole hydrochloride (Hypoxyprobe™, Hypoxyprobe Inc, Burlington, MA, USA) 90 minutes before euthanasia at day 7 or 28 post-HLI. Sections were permeabilised with 0.3% Triton X-100 (Sigma-Aldrich) in PBS, blocked with 0.5% BSA/5% goat serum for 30 minutes at RT. Slides were incubated overnight at 4°C with primary antibodies against pimonidazole adducts (1:100, Hypoxyprobe Inc) and endomucin (1:50, Invitrogen, #14-5851-82). After PBS washes, sections were incubated with Alexa Fluor^®^ 488-conjugated goat anti-rabbit (1:500, Life Technologies, #A-11008) and Alexa Fluor^®^ 647-conjugated goat anti-rat (1:500, Life Technologies #A-21246) secondary antibodies for 2 hours at RT. Nuclei were counterstained with DAPI-containing Fluoromount-G (Southern Biotech). Imaging was performed using a Zeiss LSM880 confocal microscope. Hypoxic area quantification was performed using ImageJ (NIH).

#### Vascular and lipid content analysis

Bone sections were permeabilised and blocked as above, then incubated with primary antibodies against endomucin (1:50, Invitrogen, #14-5851-82) diluted in blocking solution overnight at 4°C. After PBS washes, sections were incubated with Alexa Fluor^®^ 568-conjugated goat anti-rat (1:400; Life Technologies #A-11077) for 2 hours at RT then LipidTOX™ Green neutral lipid stain (1:200; Thermo Fisher Scientific) for 30 minutes at RT. Following final PBS washes, sections were mounted with DAPI-containing Fluoromount-G (Southern Biotech). Sections were imaged at 40x magnification using a Zeiss LSM880 confocal microscope. Image analysis was performed using ImageJ (NIH). DAPI fluorescence was used to define overall architecture. For each bone marrow sample, two 200 × 200 μm regions of interest were analysed. The percentage area of endomucin stained vasculature and LipidTOX™ stained lipid (adipocyte) staining were measured. Mean sinusoidal number and diameter were defined in the two 200 × 200 μm regions of interest per image and overall average values were produced per mouse ischaemic and non-ischaemic limb.

#### BM Flow cytometry

BM was flushed from one femur and tibia using PBS with 0.5% BSA and 2mM EDTA. Cells were washed and incubated for 10 min at 4°C with CD16/32 Fc block (BD Biosciences), then incubated for a further 10 min at 4°C with anti-CD45-VioBlue (Miltenyi Biotech), anti-Ly6G-PE (Miltenyi Biotech), anti-Ly6C-APC (Miltenyi Biotech), anti-CD11b-FITC (Miltenyi Biotech) prior to washing unbound antibodies. Separate BM samples were stained for 10 min with anti-lineage cocktail-eFluor450 (eBioscience), anti-CD117-PE (Miltenyi Biotech) and anti-Sca-1-APC (Miltenyi Biotech). Flow cytometry (CytoFlex™, Beckman Coulter) was performed to acquire leucocytes based on typical light scatter properties; with further gating to define: total CD45^+^ leukocytes, CD45^+^CD11b^+^ myeloid cells, CD45^+^CD11b^+^Ly6G^hi^Ly6C^hi^ neutrophils, CD45^+^CD11b^+^Ly6G^−^Ly6C^+^ monocytes, CD45^+^CD11b^+^LY6G^−^Ly6C^hi^ inflammatory monocytes. BM HSCs were defined as Lineage-Sca-1^+^c-Kit^+^ (LSK). All populations are expressed as cells/femur. Representative imagines of the gating strategies are provided in **Supplemental Figure 1**.

#### BMDM culture and LPS stimulation

Hindlimbs were disinfected with 70% ethanol, and femurs and tibiae harvested under sterile conditions. BM was flushed with PBS, centrifuged and red blood cells lysed with buffer (Invitrogen) for 5 minutes on ice. Cells were resuspended in differentiation medium (DMEM with stable glutamine, 20% FBS, 20 μg/mL Antibiotic-Antimycotic and 10 ng/ml murine colony stimulating factor (M-CSF)). Cells were seeded at 9 × 10^6^ cells in 8 mL per 100 × 20 mm low-adherence plate and cultured at 37°C in 5% COD with medium changes every 3 days for 7 days until fully differentiated into M0 macrophages. On day 7, naïve macrophages were washed with PBS and cultured with either differentiation medium or for M1 polarisation with 100 ng/ml LPS in DMEM containing 10% low-endotoxin FBS and 20 μg/mL Antibiotic-Antimycotic for 24 hours. Cells were detached using 2 mM EDTA/PBS for 15 minutes followed by gently scraping from the plate. After blocking with CD16/32 Fc block for 10 minutes at 4°C, cells were stained with anti-CD45-VioBlue, anti-CD11b-FITC and anti-F4/80-APC for 10 minutes at 4°C, washed, and analysed by flow cytometry (CytoFlex™, Beckman Coulter) as shown in **Supplemental Figure 2**.

#### RNA isolation, sequencing and bioinformatics

Total RNA was isolated from naïve or LPS-stimulated BMDMs after 7 days of differentiation using TRI Reagent (Sigma-Aldrich) and the RNA Clean & Concentrator-5 kit (Zymo: R1012) with DNase I treatment. RNA-seq was performed by the Leeds Next Generation Sequencing Facility (Illumina NextSeq 500 platform) to acquire 75 bp single-end reads. Raw data are deposited at (https://www.ebi.ac.uk/biostudies/arrayexpress) ArrayExpress under accession number E-MTAB-15624

Raw sequencing data were assessed for quality using FastQC v0.12.0,(8) with adapter sequences removed using TrimGalore v0.6.6.(9) Reads were aligned to the mouse reference genome (GRCm39) using STAR aligner v2.7.10a,(10) and gene-level read counts generated using featureCounts v2.0.1.(11) Differential gene expression analysis was performed using DESeq2 v1.49.3 in R v4.4.2 using RStudio.(12) Genes with raw counts <10 in ≤3samples were excluded and Apeglm shrinkage was applied to reduce false positive hits from weakly expressed genes.(13) Data were transformed using the ‘vst’ function of DESeq2 prior to generating principal component plots.

#### DNA isolation, ATAC sequencing and bioinformatics

BMDM were cultured for 7 days, treated with DNase solution (Zymo: R1012) for 30 min at 37°C, detached using TrypLE Express Enzyme (Thermo), and cryopreserved in culture medium containing 50% FBS and 10% DMSO. Cryopreserved cells were sent to Active Motif Inc. for further processing including tagmentation and library preparation based on the method of Buenrostro *et al*,(14) with minor adaptations from the method of Corces *et al*.(15) Downstream data analysis were performed by Active Motif Inc., as detailed in other published work.(16) In brief, the resulting paired-end DNA libraries were sequenced using Illumina NextSeq 500 platform with a read length of 42 bp, with over 60 million reads/sample. Raw reads were aligned to the mouse genome (mm10), and DESeq2 was used to normalise and perform differential accessibility analysis.

#### Statistics

Results are expressed as mean ±□SEM. Statistical comparisons were performed using paired Student’s t-tests, when comparing the ischaemic and non-ischaemic limbs of the same mice, or unpaired Student’s t-tests when comparing mice undergoing HLI to controls. All tests were two-sided with p <0.05 considered statistically significant. Data were analysed using GraphPad Prism 10 (GraphPad Software Inc., Boston, MA). *n* denotes the number of mice per experiment.

## Results

### HLI induces BM ischaemia

To confirm that our HLI model of left femoral artery ligation acutely impairs limb perfusion, and to monitor the kinetics of perfusion recovery, we performed serial laser Doppler perfusion imaging (**Figure 1A,B**). Immediately post-operatively, left hindlimb flux was 85% lower than the sham-operated right hindlimb; this progressively recovered with time, being 40% lower than the right hindlimb after 61 days. As this method can only define the perfusion of superficial tissues, and does not directly confirm hypoxia, we injected mice with Hypoxyprobe™ (pimonidazole) prior to harvesting tibiae from the left and right limbs 7- and 28-days post HLI induction. Under low oxygen tensions (pO2 < 10mmHg), pimonidazole is reductively activated, resulting in its stable adduction with protein thiols that can be detected with immunofluorescence.(17) BM from the right tibia exhibited discrete regions of hypoxia, remote from blood vessels, in keeping with the known presence of hypoxic niches required for HSC quiescence.(18) In contrast, there was diffuse Hypoxyprobe staining in the left tibial BM 7-days post-HLI, which had normalised by 28-days post-HLI (**Figure 1C**). Quantification of this in the tibia and femur illustrates the marked increase in Hypoxyprobe staining area on day 7, although there was large regional variability and this did not reach statistical significance (**Figure 1D**). This potentially reflects the marked derangement of tissue architecture and loss of cellularity noted with DAPI staining of nuclear DNA, which we explored next.

**Figure 1:**
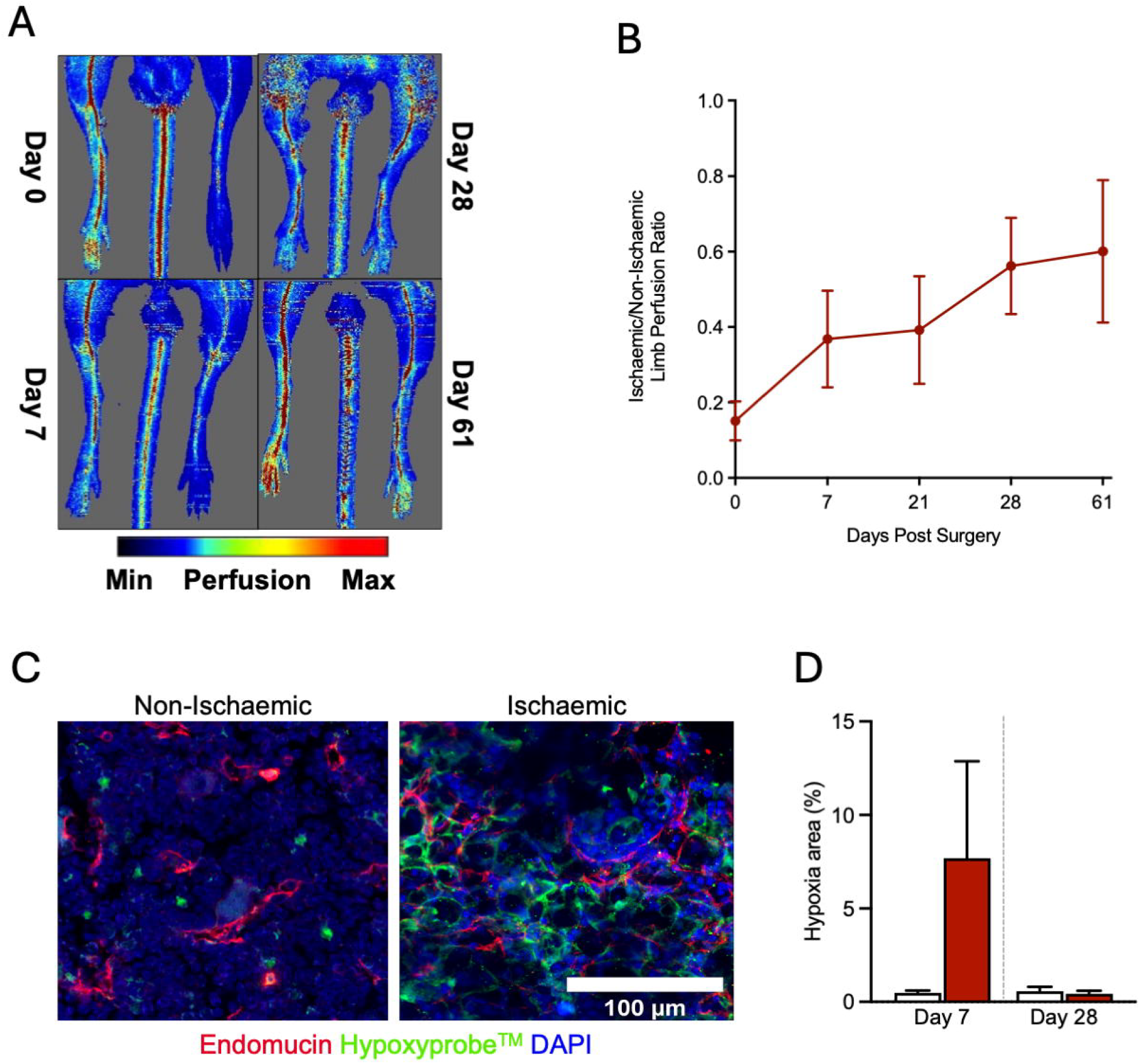
HLI surgery transiently reduces limb perfusion and induces temporary BM ischaemia. **A)** Representative images of laser Doppler flux imaging pre/post HLI. **B)** Quantification of laser Doppler data from days 0 to 61 (n=6). **C)** Representative confocal microscopy images of BM hypoxia distribution determined using Hypoxyprobe™ adducts in ischaemic and non-ischaemic tibae. Scale bar denotes 100μm; red – endomucin, green – Hypoxyprobe™, blue – DAPI-stained DNA. **D)** Quantification of Hypoxyprobe™ area in non-ischaemic (white bars) and ischaemic (red bars) BM at 7- and 28-days post-HLI (n=5,5). BM – bone marrow; HLI – hind limb ischaemia.

### HLI induces transient vascular remodelling and sustained lipid deposition

To characterise BM structural changes after HLI, we performed immunofluorescence of tibial cryosections from the ischaemic (left) and non-ischaemic (right) hindlimbs, according to the methods of Kusumbe *et al*.(7) Vasculature was defined by endomucin staining (which identifies all vessels apart from arteries), cellular content by DAPI staining, and neutral lipid content by LipidTOX™ staining. On day 7, profound abnormalities were noted in the ischaemic BM, with deranged architecture, a loss of cellularity, presence of large lipid droplets, and abundant vasculature with markedly dilated sinusoidal lumens (**Figure 2A,B**). In contrast, no morphological abnormalities were evident in the non-ischaemic BM (**Figure 2A**). However, quantification revealed a trend toward increased endomucin area and significantly decreased vascular sinusoidal area in non-ischaemic BM versus healthy BM from control mice (**Figure 2B-D**), indicating that HLI impacts BM beyond the ischaemic limb. In comparison with the non-ischaemic limb BM at day 7, the ischaemic BM had significantly greater endomucin area, more abundant vascular sinusoids, and larger sinusoidal lumen area, supporting the presence of profound vascular alterations. Moreover, lipid area was already significantly greater in the ischaemic versus non-ischaemic BM at day 7 (**Figure 2E**).

**Figure 2:**
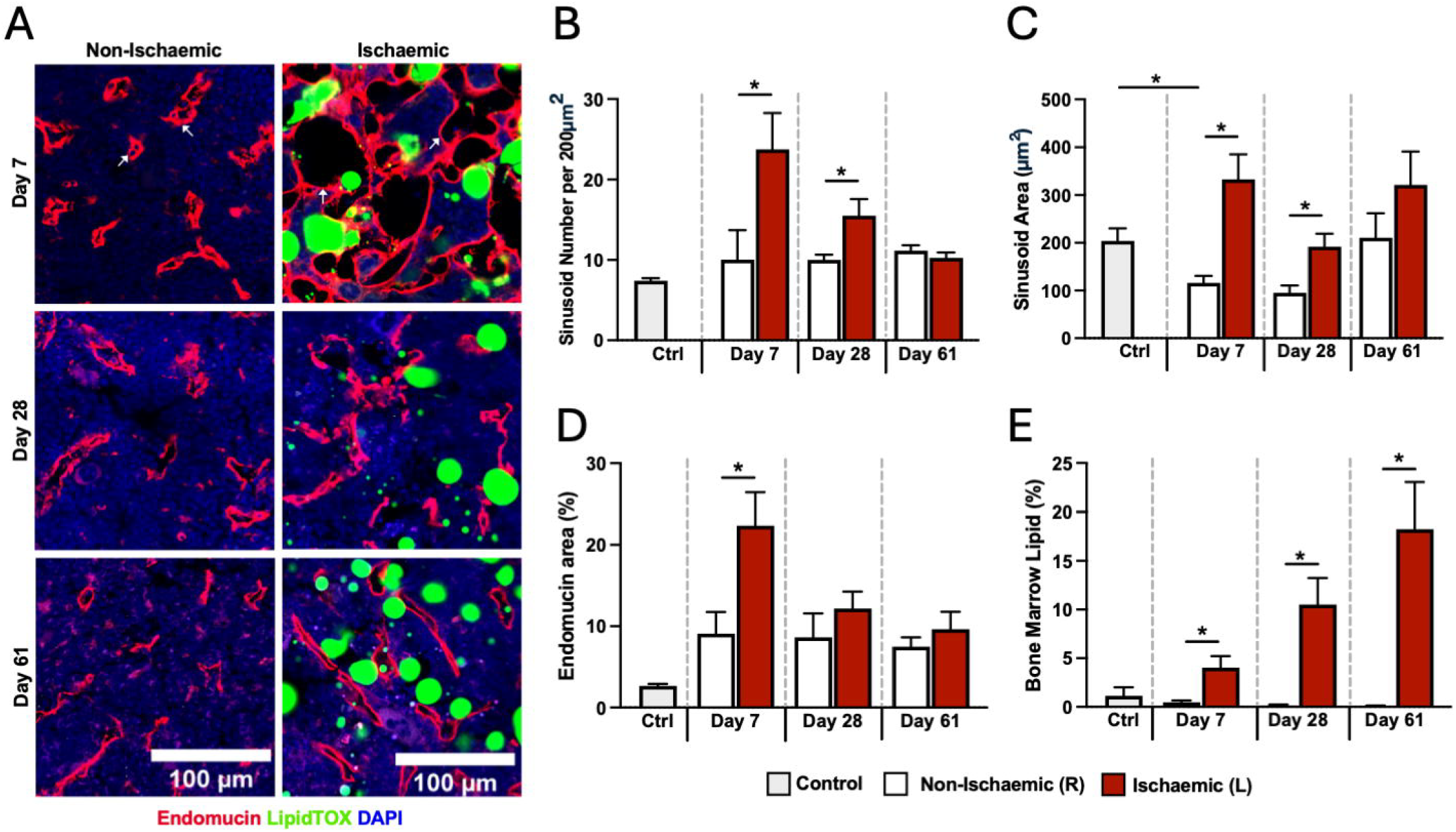
HLI induces transient and sustained remodelling of the ischaemic limb BM. **A)** Representative immunofluorescent confocal microscopy of ischaemic limb and non-ischaemic limb BM at 7-, 28- and 61-days post-HLI surgery, illustrating cell nuclei (DAPI - blue), lipid (LipidTOX – green) and vasculature (endomucin – red), with white arrows highlighting sinusoids that are profoundly disrupted at day-7 in the ischaemic BM. Scale bars denote 100μm. **B-D)** Quantification of BM vascular properties in ischaemic limb and non-ischaemic limb BM at 7-, 28- and 61-days post-HLI surgery. (n=6,6) **E)** Quantification of BM lipid content in ischaemic limb and non-ischaemic limb BM at 7-, 28- and 61-days post-HLI surgery. (n=6,6) * denotes p<0.05 in ischaemic versus non-ischaemic BM.

Further analyses at day 28 post-HLI revealed partial normalisation of vascular abnormalities in the ischaemic limb BM, which had fully normalised by day 61 post-HLI. However, there was a stepwise increase in lipid area in the ischaemic BM during this time, which accounted for over 15% of the BM area by day 61 (**Figure 2B-E**). Hence, whilst the profound vascular abnormalities induced by HLI partially normalise over time, there appears to be a progressive and sustained increase in lipid deposition, suggesting long-standing structural abnormalities develop after BM ischaemia.

### HLI alters haematopoiesis in ischaemic and non-ischaemic hind limbs

Given the primary role of the BM in haematopoiesis, and the apparently lower cellularity noted on immunofluorescence after HLI, we next characterised the haematopoietic cellular content of the ischaemic and non-ischaemic BM using flow cytometry. (**Figure 3**) On day 7 post-HLI, we observed no difference between ischaemic and non-ischaemic limb BM abundance of: total cells; CD45^+^ leukocytes; CD45^+^CD11b^+^ myeloid cells; CD45^+^CD11b^+^Ly6G^hi^Ly6C^hi^ neutrophils; CD45^+^CD11b^+^Ly6G^−^Ly6C^+^ monocytes; CD45^+^CD11b^+^Ly6G^−^Ly6C^hi^ inflammatory monocytes; or lineage^−^Sca-1^+^c-Kit^+^ HSCs. However, in comparison with control mice, both the ischaemic and non-ischaemic BM exhibited significantly higher monocyte and inflammatory monocyte numbers. These changes persisted at day 61 post-HLI in the non-ischaemic BM. On day 61 post-HLI, all quantified cell lineages other than HSCs were significantly more abundant in the non-ischaemic than the ischaemic BM, with the ischaemic BM counts more closely resembling those of control mice. However, the abundance of HSCs was markedly reduced versus control mice in both the ischaemic and non-ischaemic BM. These data indicate sustained abnormalities in BM cellular composition after HLI, particularly of the non-ischaemic BM, highlighting enduring systemic effects of HLI on haematopoiesis, even after resolution of abnormal BM hypoxia.

**Figure 3:**
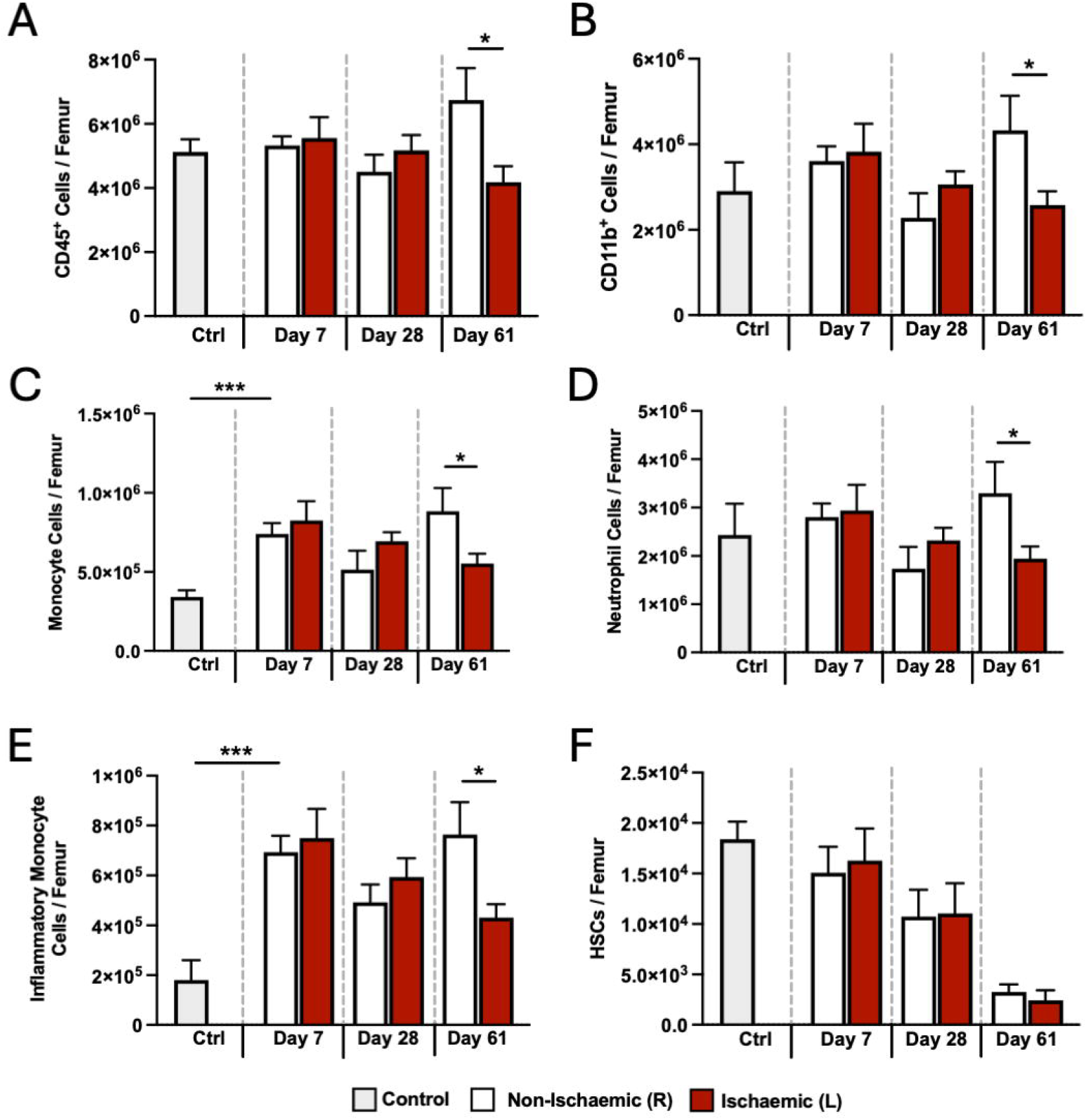
HLI induces transient and sustained alterations to haematopoiesis in the ischaemic and non-ischaemic BM. Flow cytometry quantification of **A)** total CD45^+^ leukocytes, **B)** CD45^+^CD11b^+^ myeloid cells, **C)** CD45^+^CD11b^+^Ly6G^hi^Ly6C^hi^ neutrophils, **D)** CD45^+^CD11b^+^Ly6G^−^Ly6C^+^ monocytes, **E)** CD45^+^CD11b^+^LY6G^−^Ly6C^hi^ inflammatory monocytes, and **F)** Lineage^−^Sca-1^+^c-Kit^+^ HSCs from BM of control mice, and the ischaemic or non-ischaemic limb BM of mice 7-, 28- and 61-days post-HLI. BM – bone marrow; HLI – hind limb ischaemia; HSCs - haematopoietic stem cells. (n=6,6) * denotes p<0.05 in ischaemic versus non-ischaemic BM and *** denotes p<0.05 in control versus non-ischaemic BM.

### HLI alters BMDM gene expression

Given the prominent increase in BM monocyte count post-HLI, we next explored the function of BMDMs. To derive these, BM was separately collected from the ischaemic and non-ischaemic limbs 7 days post-HLI, along with BM from control mice. BMDMs were cultured for 7 days in normoxic conditions, prior to confirming uniform expression of CD45, CD11b and F4/80 in all samples using flow cytometry (**Supplemental Figure 2**). A further 24-hour exposure to standard culture media was used to generate naïve (non-polarised or M0) BMDMs, whereas standard media with 100ng/ml LPS was used to induce polarisation of BMDMs to an M1-like state, prior to performing bulk RNA-seq.

A principal component (PC) analysis plot revealed clear separation of naïve and LPS-stimulated samples on PC1, which accounted for 72% of the transcriptional variance (**Figure 4A**). Appropriate augmentation of canonical LPS-responsive genes, such as Tnf, Il1b and Nos2 (all p<1×10^−50^), was seen amongst 6,960 DEGs in control-LPS versus control-naïve BMDMs (**Supplemental Table 1**). PC2, which accounted for 20% of transcriptional variance, appeared to reflect HLI status, with clear separation of naïve-BMDMs from control, non-ischaemic and ischaemic BM. When comparing naïve ischaemic to naïve control BMDMs, there were 3,635 DEGs (**Supplemental Table 2**), functional enrichment of which resulted in Gene Ontology Biological Process (GO-BP) terms including ‘innate immune response’ (p=4.6×10^−45^) and cholesterol metabolic process (p=2.8×10^−11^)(**Supplemental Table 3**). Notably, there was no differential expression of genes indicating M2 polarisation, such as Arg1, Mrc1 or Cd163.(19) Comparison of naïve non-ischaemic to naïve control yielded 1,771 DEGs (**Supplemental Table 4**), with functional enrichment highlighting GO-BP terms ‘actin filament-based process’ (p=3.3×10^−28^) and response to interferon-beta (p=2.5×10^−21^)(**Supplemental Table 5**). Cross-referencing these DEGs revealed 1,048 shared by the ischaemic and non-ischaemic BMDMs, with 723 unique to the non-ischaemic BMDMs and 2,599 unique to ischaemic BMDMs (**Figure 4B**). These data show that HLI has substantial local and systemic effects on naïve BMDMs, with both shared and distinct adaptations occurring in the ischaemic and non-ischaemic limbs.

**Figure 4:**
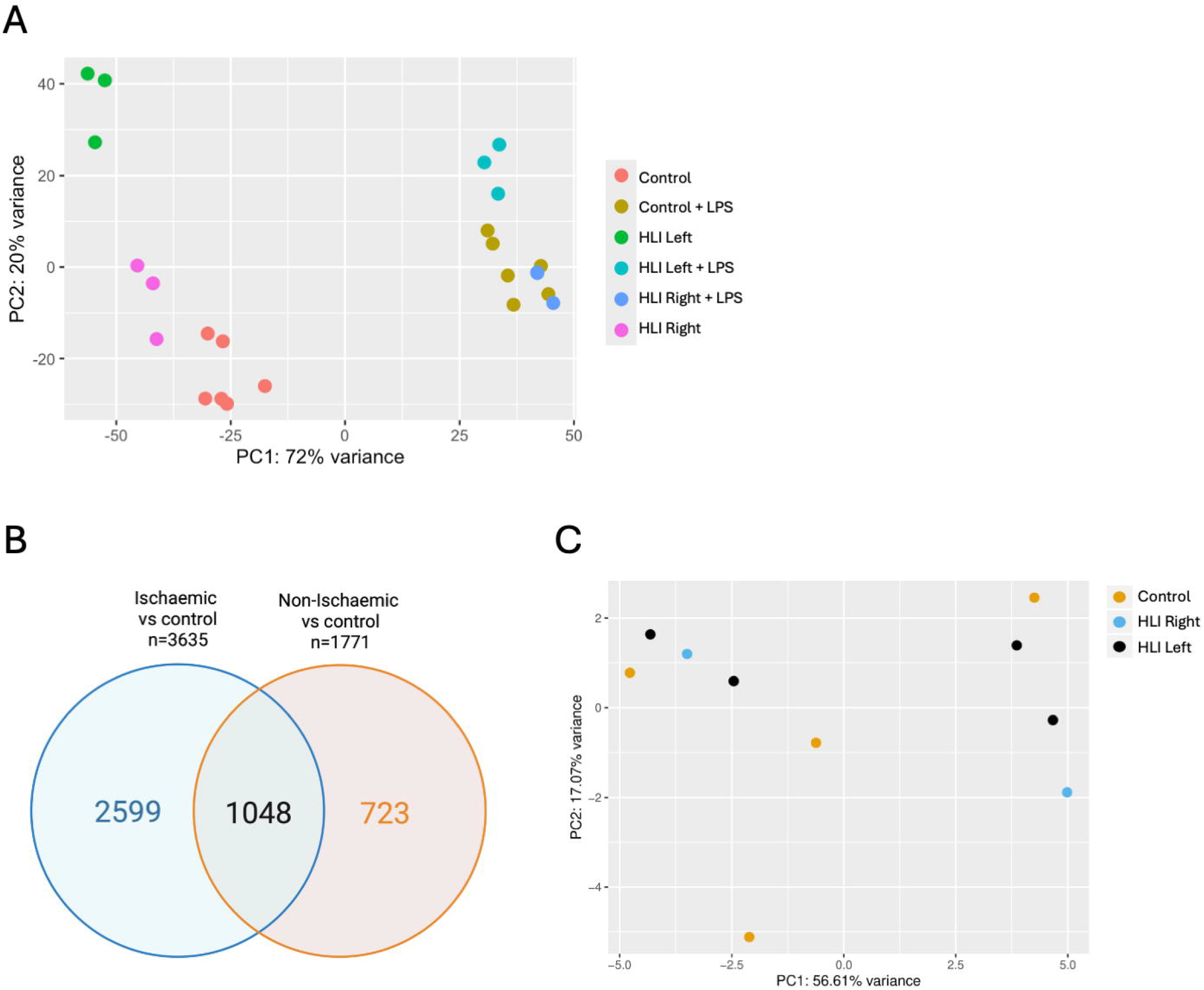
HLI induces transcriptomic alterations in ischaemic and non-ischaemic limb-derived BMDMs without marked changes in chromatin accessibility. **A)** Principal component analysis of RNA-sequencing data from BMDMs derived from control mice and the ischaemic or non-ischaemic limbs of mice 7-days post-HLI surgery, both in their naïve state and after LPS stimulation for 24-hours. (control n=6, control LPS n=6, all other conditions n=3) **B)** Venn diagram illustrating number of DEGs in comparison of naïve ischaemic limb BMDMs versus naïve control BMDMs and naïve non-ischaemic limb BMDMs versus naïve control BMDMs. **C)** Principal component analysis of ATAC-sequencing chromatin accessibility data from BMDMs derived from control mice and the ischaemic or non-ischaemic limbs of mice 7-days post-HLI surgery, all in their naïve state. (control n=4, sham limb HLI R n=2, ischaemic HLI L n=4) BMDMs – bone marrow derived macrophages; DEGs – differentially expressed genes; HLI – hind limb ischaemia.

Next, we asked whether HLI influenced the response of BMDMs to LPS. As described above, PC2 reflected HLI status in naïve BMDMs, but this pattern was less apparent in the context of LPS-stimulation, with overlapping of the control-LPS and non-ischaemic-LPS clusters. However, there remained distinct clustering of the ischaemic-LPS samples, indicating that the response to LPS differed in the context of HLI. Indeed, 411 DEGs were found between ischaemic-LPS versus control-LPS BMDMs (**Supplemental Table 6**), with functional enrichment analysis identifying terms including ‘actin polymerization-dependent cell motility’ and ‘sequestering of BMP from receptor via BMP binding’ (both p=4.3×10^−5^; **Supplemental Table 7**). To complement this, we also defined genes with statistically significant interaction between ischaemic limb (versus control) and LPS exposure (versus naïve). This revealed 1,705 hits, 687 with a positive interaction (i.e. LPS induced a greater increment in gene expression in ischaemic than control BMDMs) and 1,018 with a negative interaction (**Supplemental Table 8**). Functional enrichment analysis of genes with positive interaction yielded terms including ‘endocytosis’ (p=6.2×10^−21^) and ‘lamellipodium organization’ (p=2.7×10^−7^) (**Supplemental Table 9**), whilst those with negative interaction yielded terms including ‘actomyosin structure organization’ (p=7.5×10^−9^) and ‘regulation of transforming growth factor beta receptor signaling pathway’ (p=1.4×10^−8^; **Supplemental Table 10**). Hence, beyond its impact on naïve BMDMs, HLI also influences the response of ischaemic limb BMDMs to LPS.

Finally, given that HLI induced transcriptional differences in BMDMs after 7 days cultures in standardised normoxic conditions, we asked if there was an epigenetic basis for this, akin to that demonstrated in the immune training literature.(20) To explore this, we performed ATAC-sequencing to define differential chromatin accessibility in naïve ischaemic versus naïve control BMDMs (n=4 per group, with a further 2 non-ischaemic naïve samples analysed for further context in PC analyses). Contrary to our hypothesis, there was no clustering of samples according to group (**Figure 4C**), with only 2 genomic regions demonstrating differential chromatin accessibility between ischaemic and control groups (**Supplemental Tables 11 and 12**). Of these, only one was within 10Kb of a gene (Atp8a2), but this was not noted in our earlier list of 3,635 DEGs between ischaemic naïve and control naïve BMDMs (**Supplemental Table 2**), nor the 411 DEGs between ischaemic-LPS versus control-LPS BMDMs (**Supplemental Table 6**). Whilst not excluding the contribution of epigenetic modifications to the transcriptomic effects of HLI, these data do not support the presence of marked differences in chromatic accessibility.

## Discussion

Whilst a small number of papers have highlighted the impact of limb ischaemia on bone marrow-derived myeloid cells, our work provides significant contributions to the literature on BM ischaemia. First, we show that HLI induces BM hypoxia that is associated with profound, but transient vascular remodelling in the ischaemic limb. Additionally, there is a sustained accumulation of BM lipid in the ischaemic limb, which increased even between the first- and second-month post-HLI. This was associated with increased abundance of monocytes in both the ischaemic and non-ischaemic limb BM 7-days post-HLI, with this becoming more prominent in the non-ischaemic BM after two months, indicating important systemic effects of HLI. Our second important finding was that HLI induces transcriptional abnormalities in macrophages derived from both the ischaemic and non-ischaemic limb BM, despite their culture in normoxia for 7-days. However, macrophages from the ischaemic limb exhibited much more profound transcriptional abnormalities and also responded differentially to stimulation with LPS, suggesting some effects of HLI are localised rather than systemic. Finally, our analyses did not demonstrate major epigenetic modifications in these macrophages using ATAC-seq, which may suggest other mechanisms underpin the major transcriptomic changes that we observed after HLI.

### Structural responses to BM ischaemia

Our data show that murine HLI transiently increases BM hypoxia, which is associated with a marked increase in vascularity and loss of normal sinusoidal architecture that broadly normalises over months. Notably, healthy BM contains hypoxic regions that are thought to provide a niche for HSC, and our data found evidence of such regions in the non-ischaemic BM. Interestingly, the transcription factors Yap1 and Taz have been shown to limit BM angiogenesis in response to hypoxia, which is thought to be important in maintaining hypoxic environment in HSC niche.(21) How this physiological adaptation is modified during ischaemia is an important question that warrants further exploration.

BM lipid is stored in specialised BM adipocytes and its content tends to increase with age and in the context of many diseases.(22, 23) We observed marked increases in BM lipid after HLI and, in contrast to transient vascular perturbations, these were sustained for the duration of our study. Whether this is a hypoxia-dependent phenomenon is not certain, although femoral head magnetic resonance imaging of patients with non-atherosclerotic avascular necrosis corroborates lipid accumulation in ischaemic human BM.(24) Moreover, histology of femoral BM from patients with CLTI and T2DM shows fatty replacement of the BM, along with a marked reduction in HSCs,(25) in keeping with our data. Whether BM adipocytes accumulating post-HLI are functionally distinct from those in non-ischaemic BM remains to be explored. Research by Ferland-McCollough *et al* has shown pro-inflammatory adipocyte accumulation in the BM of people with diabetes;(26) pro-inflammatory BM adipocytes have been shown to actively promote loss of HSCs,(27) which could explain why we saw progressive adipocyte accumulation associated with HSC loss.

### Haematopoiesis post-HLI

Despite marked changes in BM vasculature and lipid content post-HLI, we observed no differences between ischaemic and non-ischaemic limb BM (nor versus control BM) in the abundance of total CD45+ leukocytes or CD11b+ myeloid cells in the first month. Whilst there were no differences between the ischaemic and non-ischaemic limb BM monocyte (and inflammatory monocyte) abundance, this was markedly higher in both limbs versus control mice at day 7 and remained higher in the non-ischaemic limb. This was paralleled by a stepwise decline in lineage^−^Sca-1^+^c-Kit^+^ HSCs in both limbs, versus control mice, in the first and second months after HLI. This suggests important changes in haematopoiesis, that may be more reflective of the systemic response to HLI than the local implications of BM ischaemia, given the similar or more pronounced changes in the non-ischaemic limb. Interestingly, other research has shown that HLI induces inflammation within contralateral limb muscle, further supporting the notion of effects.(28)

The functional implications of altered haematopoiesis in the ischaemic and non-ischaemic BM are also important, for example in systemic inflammatory responses. Our macrophage RNA-seq studies found clear transcriptional differences between control macrophages and those from mice post-HLI, especially macrophages derived from the ischaemic limb BM. In the context of stimulation with LPS, ischaemic limb macrophages remained transcriptionally difference from control macrophages. Data from Falero-Diaz *et al* revealed that 24-hours of hind-limb ischaemia-reperfusion injury leads to a sustained change in the transcriptome and lipidome of ischaemic limb BM-monocytes.(3) Functionally, these monocytes were more able to promote ischaemic muscle angiogenesis and blood flow recovery post-HLI when adoptively transferred. Their subsequent research revealed ischaemia-reperfusion to prime BM-macrophages to an anti-inflammatory M2-like phenotype.(4) Notably, we did not observe differential expression of canonical M2 marker genes (e.g. Arg1, Mrc1), and whilst Falero-Diaz found lower Dhcr24 expression to be important to their observations, we found expression of this to be significantly higher in ischaemic limb macrophages. These differences may reflect the distinction between transient ischaemia-reperfusion and sustained limb ischaemia, but further work is needed to understand this.

Coppin *et al* demonstrated that sustained HLI in atherosclerosis prone ApoE^−/−^ mice induced pro-inflammatory myeloid cell egress from the BM, associated with increased atherogenesis.(5) Using single cell RNA-sequencing (RNA-seq), they found perturbations in inflammatory gene expression in all myeloid cell populations, along with altered gene expression in their progenitors and HSCs, although do not specifically state if this refers to BM from the ischaemic and/or non-ischaemic limbs. Their data is valuable in characterising responses in multiple cell populations *in vivo*, and this approach would be useful in extending our observations on altered LPS responses. Interestingly, Coppin *et al* found evidence for altered epigenetic modifications associated with differential expression of Il1r and Il3rb, in contrast to our ATAC-seq data. This difference may reflect their focussed analysis on specific chromatin modifications (rather than chromatin accessibility), greater sample size, and the reduced statistical penalty from focussing on 2 genes. An epigenetic basis for our observed differences remains credible and a larger scale ATAC-seq analysis would help to address this in future.

## Limitations

It is also important to acknowledge and discuss the limitations of our study. First, we only present from a murine model and this does not replicate the complexity of human CLTI. Whilst our data help to explore the implications of bone marrow ischaemia locally and systemically, human studies will be essential to define the translational potential of these data in terms of biomarker derivation and therapeutic relevance. Second, we do not explore the mechanistic basis for our observations which are likely to be complex and involve multiple cell lineages, hence requiring a large programme of research. Many important questions remain about the basis of transient and sustained changes in the BM, including for haematopoiesis. Third, the sample size of some experiments, especially ATAC-seq, is modest and increases the risk of false negative findings as discussed above. Moreover, our characterisation of haematopoiesis was focussed and so many questions remain about the implications of HLI for myeloid responses to stimuli beyond LPS and also regarding the impact on other lineages (e.g. erythroid).

## Conclusions

Our study adds to the emerging data defining how murine limb ischaemia impacts BM structure and function. We show that HLI induces transient BM hypoxia, associated with temporary vascular remodelling and sustained accumulation of BM lipid. These changes are associated with altered abundance and function of leukocytes in not only the ischaemic limb, but also the contralateral non-ischaemic limb. Moreover, macrophages derived from the ischaemic BM showed altered responses to LPS, suggesting implications for pathogen responses. Future work should explore these phenomena in people with CLTI with a view to understanding the systemic implications of this disease, developing biomarkers of BM-ischaemia and identifying therapeutic opportunities.

## Acknowledgements

This work was funded by the British Heart Foundation (FS/18/61/34182). Equipment used in this work was provided by the University of Leeds Bioimaging and Flow Cytometry facility which is supported by funding from the Wellcome Trust (WT104918MA) and Biotechnology and Biological Sciences Research Council (BBSRC BB/R000352/1).

## Supplemental data

**Supplemental Figure 1: Flow cytometry profile of bone marrow haematopoietic cells**. Representative scatterplots and histograms showing gating strategy to define total CD45^+^ leukocytes, CD45^+^CD11b^+^ myeloid cells, CD45^+^CD11b^+^Ly6G^hi^Ly6C^hi^ neutrophils, CD45^+^CD11b^+^Ly6G^−^Ly6C^+^ monocytes, CD45^+^CD11b^+^LY6G^−^Ly6C^hi^ inflammatory monocytes, and Lineage^−^Sca-1^+^c-Kit^+^ HSCs in bone marrow.

**Supplemental Figure 2: Flow cytometry profile of cultured BMDMs**. Representative histogram and scatterplot confirming almost uniform expression of CD45, CD11b and F4/80 in bone marrow-derived macrophages (BMBMs) cultured *in vitro* for 7 days.

**Supplemental Table 1: Differentially expressed genes in control BMDMs exposed to LPS versus naïve control BMDMs**. Data presented in .csv format

**Supplemental Table 2: Differentially expressed genes in naïve ischaemic BMDMs versus naïve control BMDMs**. Data presented in .csv format

**Supplemental Table 3: Gene Ontology terms enriched in differentially expressed genes from naïve ischaemic BMDMs versus naïve control BMDMs**. Data presented in .csv format

**Supplemental Table 4: Differentially expressed genes in naïve non-ischaemic BMDMs versus naïve control BMDMs**. Data presented in .csv format

**Supplemental Table 5: Gene Ontology terms enriched in differentially expressed genes from naïve non-ischaemic BMDMs versus naïve control BMDMs**. Data presented in .csv format

**Supplemental Table 6: Differentially expressed genes in LPS-stimulated ischaemic BMDMs versus LPS-stimulated control BMDMs**. Data presented in .csv format

**Supplemental Table 7: Gene Ontology terms enriched in differentially expressed genes from LPS-stimulated ischaemic BMDMs versus LPS-stimulated control BMDMs**. Data presented in .csv format

**Supplemental Table 8: Genes with expression exhibiting statistically significant interaction between ischaemic limb (versus control) and LPS exposure (versus naïve)**. Data presented in .csv format

**Supplemental Table 9: Gene Ontology terms enriched in genes with expression exhibiting statistically significant positive interaction between ischaemic limb (versus control) and LPS exposure (versus naïve)**. Data presented in .csv format

**Supplemental Table 10: Gene Ontology terms enriched in genes with expression exhibiting statistically significant negative interaction between ischaemic limb (versus control) and LPS exposure (versus naïve)**. Data presented in .csv format

**Supplemental Table 11: Differential chromatin accessibility between naïve ischaemic BMDMs versus naïve control BMDMs**. Data presented for hits with false discovery rate-adjusted p<0.05 in .xlsx format

**Supplemental Table 12: Differential chromatin accessibility between naïve ischaemic BMDMs, naïve non-ischaemic BMDMs and naïve control BMDMs**. Data presented for all genes, irrespective of statistical significance, in .xlsx format

