## Supplementary figures and images for "Murine limb ischaemia induces structural and functional remodelling of local and distant bone marrow"

### Supplemental Figure 1

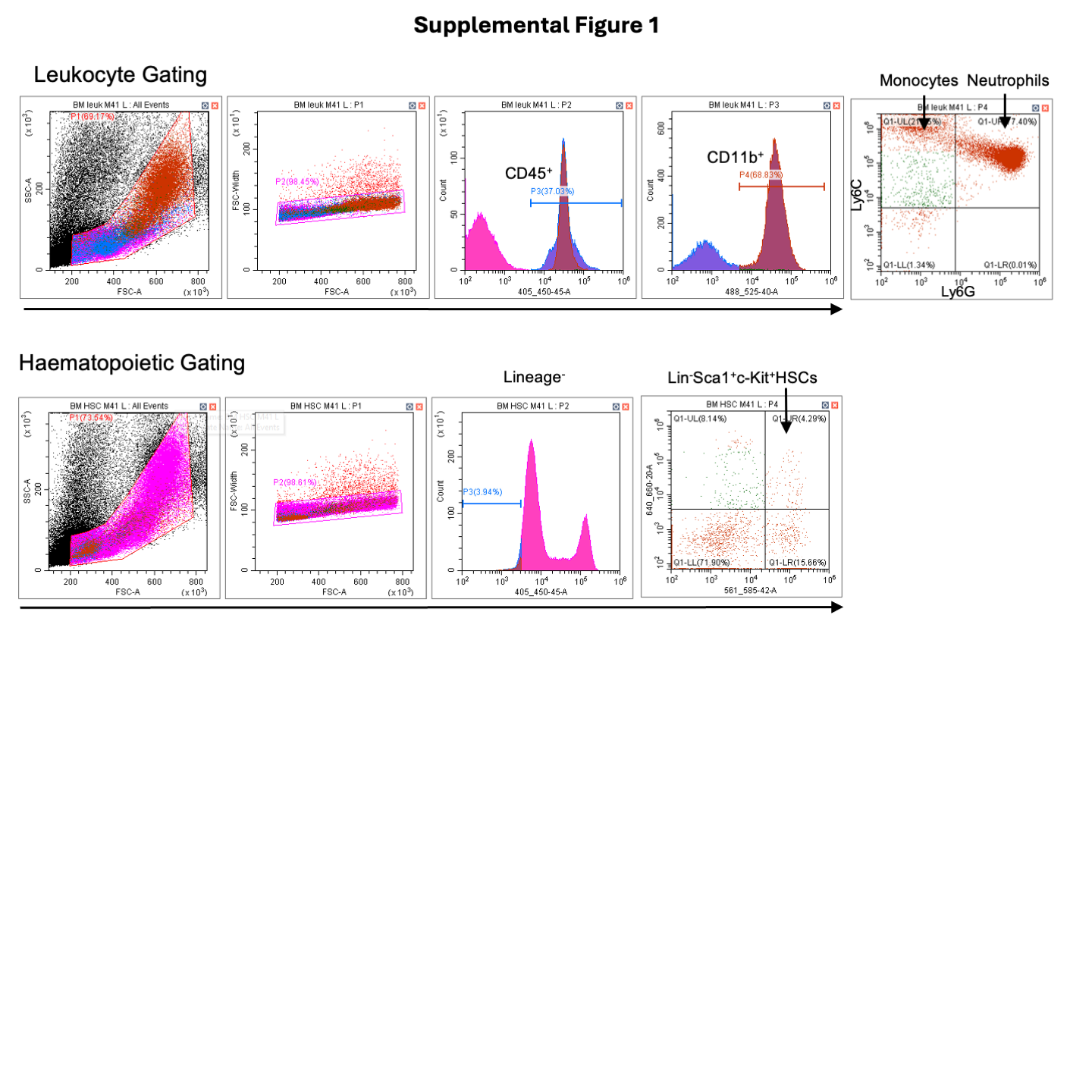

### Supplemental Figure 2

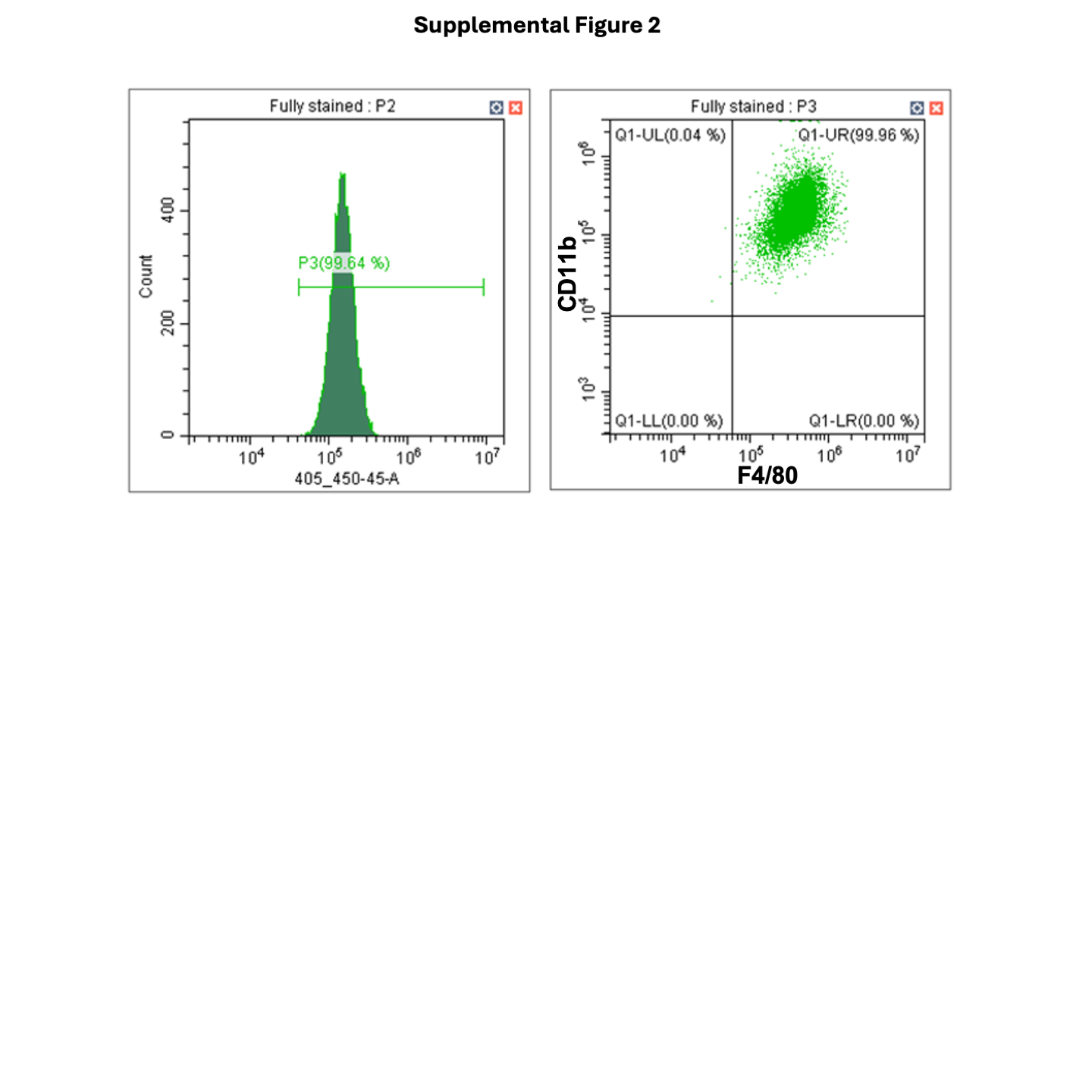
